# Improving an rRNA depletion protocol with statistical design of experiments

**DOI:** 10.1101/2022.03.10.483821

**Authors:** Benjamin M. David, Paul A. Jensen

## Abstract

rRNA depletion is the most expensive step in prokaryotic RNA-seq library preparation. rRNA is so abundant that small increases in depletion efficiency lead to large changes in mRNA sequencing coverage. A variety of commercial and home-made methods exist to lower the cost or increase the efficiency of rRNA removal. Many of these techniques are suboptimal when applied to new species of bacteria or when the protocol or reagents need to be changed. Re-optimizing a protocol by trial-and-error is an expensive and laborious process. Systematic frameworks like the statistical design of experiments (DOE) can improve processes by exploring the quantitative relationship between multiple factors. DOE allows experimenters to find factor interactions that may not be apparent when factors are studied in isolation.

We used DOE to optimize an rRNA depletion protocol by updating reagents and identifying factors that maximize rRNA removal and minimize cost. The optimized protocol more efficiently removes rRNA, uses fewer reagents, and is less expensive than the original protocol. Our optimization required only 17 experiments and identified two significant interactions among three factors. Overall, our approach demonstrates the utility of a rational, DOE framework for improving complex protocols.

## 1 Introduction

RNA-seq uses high-throughput DNA sequencing to measure mRNA transcript abundance and quantify gene expression. Over 80% of the total RNA in a prokaryotic cell is rRNA that must be removed from the sample during RNA-seq library preparation to minimize the sequencing depth required to measure mRNA abundance [1]. Removing rRNA is the largest single expense when preparing an RNA-seq library, accounting for over half of the total cost. A variety of methods—both commercial and home-made—have been developed to lower cost, improve convenience, or increase the efficiency of rRNA removal [2]–[6].

Beginning in the 1950’s, the field of quality engineering discovered that nearly any process can be improved through the statistical design of experiments (DOE) [7], [8]. Even processes that operate well can benefit from a quantitative exploration of the relationship between process variables. Frameworks like Response Surface Methodology (RSM) allow experimenters to vary multiple factors simultaneously and search for better operating conditions using a mathematical model [9]. More recently, DOE and RSM have been used to improve molecular biology protocols [10], [11].

RSM can improve molecular biology protocols in three ways. First, the original protocol developers may have stopped tuning the protocol when it worked well enough for their needs. RSM can find changes to the protocols that may not have been considered by the developers. Second, reagents used in the original protocol may not be available later or might be manufactured in a different way. RSM can adjust the protocols to account for changing reagents, equipment, or techniques. Finally, the protocol developers may not have considered the effects of some factors. Analyzing all factors in a protocol leads to a combinatorial explosion of variants, so scientists often focus on only a few factors or ignore interactions between factors. RSM and statistical DOE allow developers to efficiently measure combinatorial effects with relatively few experiments.

We used RSM and DOE to optimize a previously developed rRNA depletion protocol that uses complementary RNA probes labeled with biotin to remove hybridized rRNA with streptavadin-coated magnetic beads [2]. Compared to commercially available alternatives, this strategy is both less expensive and automation friendly, which makes it ideal for large-scale transcriptomic experiments [4], [6]. The protocol was originally developed to remove rRNA from total RNA isolated from environmental communities before RNA-seq. The protocol designers optimized rRNA depletion efficiency by designing probes that target rRNAs from multiple taxa in an environmental community. In this work, we optimized the rRNA depletion protocol by using updated reagents and finding reagent stoichiometry settings that maximize depletion efficiency and minimize cost in a single bacterial species.

## 2 Results

### 2.1 rRNA removal by hybridization

Figure 1A depicts the rRNA removal protocol described by Stewart, *et al*. [2]. The 16s and 23s rRNA genes are amplified by PCR with a T7 promoter added to the reverse primer. In vitro transcription (IVT) creates RNA probes that are complementary to the 16s and 23s rRNAs. The IVT reaction includes biotin-labeled cytosine and uracil ribonucleotides, adding biotin along the backbone of each probe. The antisense probes are mixed with the total RNA and hybridize to the rRNA. The hybrids bind to streptavidin coated magnetic beads and are separated from the other RNA. An RNA-seq library is prepared from the rRNA depleted samples.

**Figure 1:**
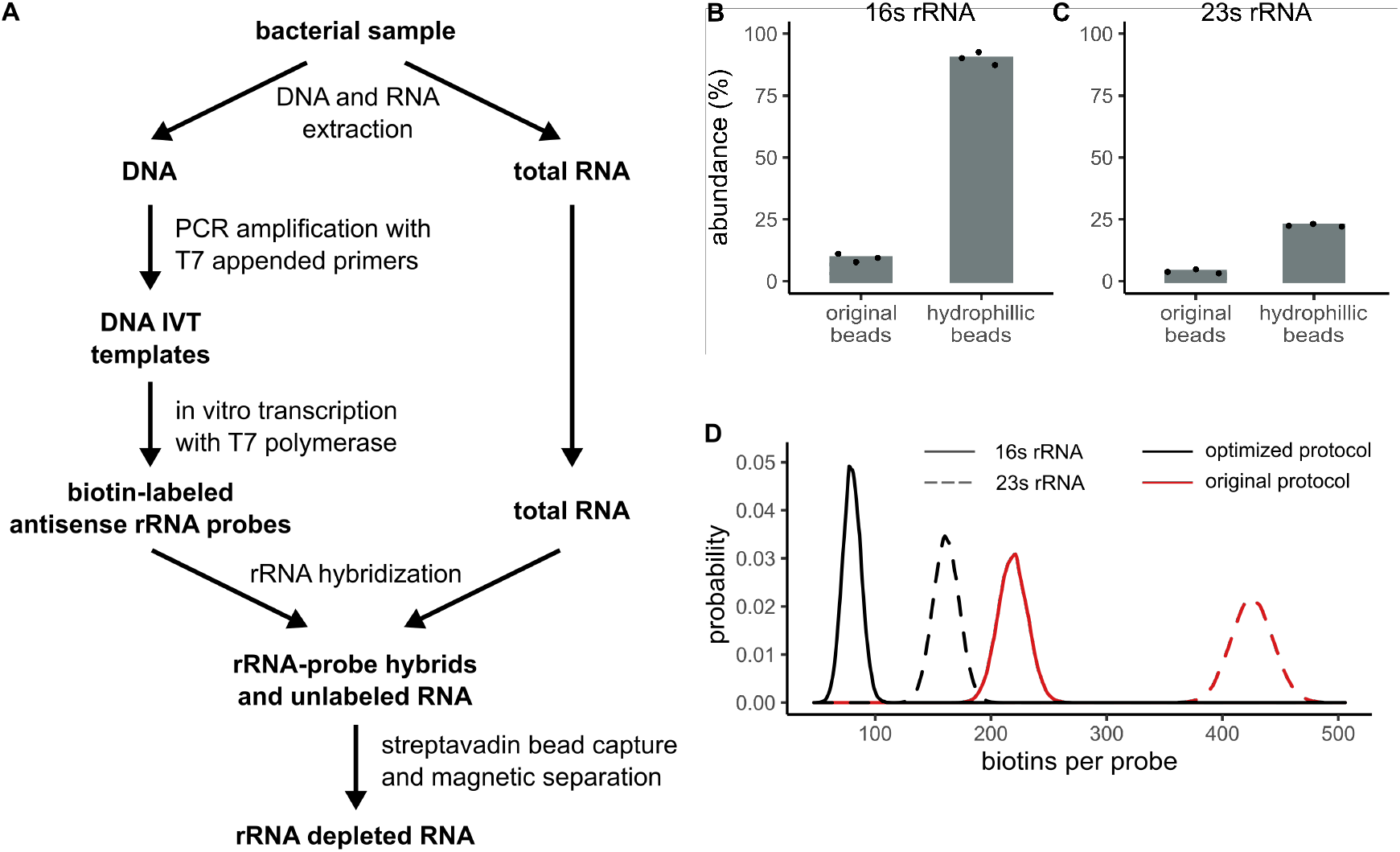
Biotin-labeled antisense rRNA probes remove rRNA from total RNA. **A**. Genomic DNA and total RNA are extracted from a bacterial sample. rRNA genes are amplified by PCR with a T7 promoter sequence included in the reverse primer. Antisense rRNA probes are synthesized in an *in vitro* transcription reaction that includes biotin-labeled nucleotides. The rRNA probes are hybridized to rRNA in the total RNA and captured by strepavadin-coated magnetic beads. **B**,**C**. replacing the original streptavadin magnetic beads with the hydrophillic variant increases the abundance of rRNA and leftover probe as measured by qPCR. **D**. Antisense rRNA probes in the original protocol contain biotin-labeled CTP and UTP. Our optimized protocol reduces the overall biotin concentration by removing biotin-labled CTP and decreasing the concentration of UTP.

Prior to optimization by DOE and RSM, we exchanged the original streptavadin magnetic beads for a new hydrophillic variant. The hydrophillic variant is recommended for use with nucleic acids because it exhibits a lower rate of non-specific nucleic acid binding; however, the hydrophillic beads have a lower overall binding capacity than the original beads [12](NEB, Personal Communication). We tested effect of switching the streptavadin bead type by depleting rRNA from total RNA samples from the bacterium *Streptococcus mutans* and measuring the amount of rRNA and probe remaining by qPCR. Following the original protocol and using the original magnetic beads, we observed an rRNA and probe abundance of 9.7% and 3.9% for the 16s and 23s rRNAs, respectively (Figure 1B,C). (The abundance is relative to the concentration of rRNA in the samples before depletion.) As expected, when using the hydrophillic beads, the measured abundance increased to 90.3% and 22.5%. Because our assay measures both leftover probe and rRNA, these results suggested that excess probe remained in the samples.

We hypothesized that probes remained in the sample because the streptavidin on the hydrophillic beads was saturated with biotin. The original protocol recommends adding 800 ng each of the 16s and 23s probes to 400 ng of total RNA, creating a 2-fold excess of probe to rRNA. Additionally, 50% of the UTPs in the IVT reaction had biotin labels, adding an average of 219 and 425 biotins per molecule of 16s and 23s probe, respectively. (To limit the number of reagents, we chose to not include CTP and double the concentration of UTP from 25% to 50% to match the original protocol (Supplementary Material 3).) To reduce the number of biotins per probe in our optimized protocol, we used only 20% of labeled UTPs. The lower concentration adds an average of 80 biotins per molecule in the shorter 16s probe with less than 1 in 5×10^38^ probes having no biotin modification (Figure 1D). Unfortunately, decreasing the number of biotins did not improve the amount of probe captured by the beads, and much of the probe remained in the sample. However, decreasing the concentration of labeled UTP and removing the CTP reduced the cost of the probe synthesis reaction by 55% (Supplemental Material 4.2).

Varying biotin concentration alone failed to reproduce the results in the original manuscript. Rather than testing additional factors one at a time, we switched to RSM to identify factors and interactions that improve rRNA depletion.

### 2.2 RSM Optimization

Optimization by RSM has three stages (Figure 2A). First, an experimental design is selected with each experiment (a *run*) having a unique combination of settings for all factors. Varying factors simultaneously makes RSM designs efficient, and the design is chosen so the effects of individual factors and their interactions can be estimated. Second, the experiments are carried out and the *response*—in our case the abundance of rRNA and probe measured by qPCR—is recorded for each run. A linear model is fit to the experimental data to estimate main (first order) effects, two way interactions, and pure quadratic effects for each factor. Finally, the model directs the search for new factor settings that optimize the response. The new factor settings are tested, and the experimenter can repeat the RSM process to further improve the process in a new design space.

**Figure 2:**
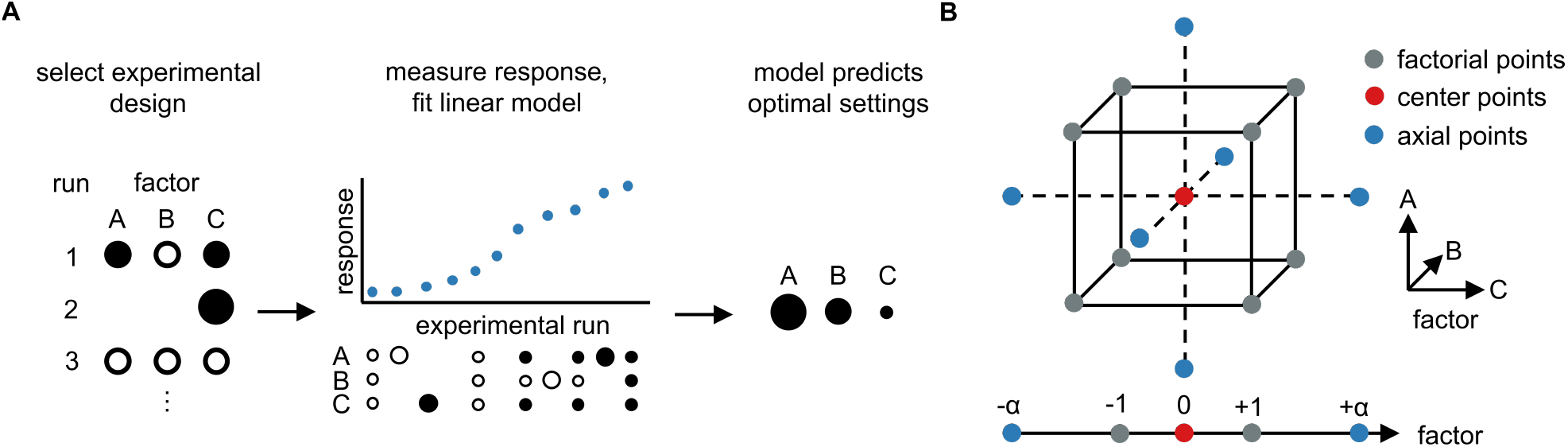
Response surface methodology (RSM) is used for process optimization. **A**. RSM is performed in three stages. First, a multi-factorial experimental design is selected. Second, the experiment is carried out, the response is measured, and a mathematical model is fit to the response surface. Third, the model guides the search for optimal settings. **B**. A 3-factor rotatable central composite design (CCD) was chosen to assess first order, two-way interaction, and quadratic effects. The CCD has a factorial core to map first order and two-way interaction effects, and center and axial points to measure quadratic effects. For a rotatable CCD, the coded level for the axial points (*α*) is set at 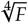, where *F* is the number of factorial points.

Compared to other DOE methodologies, RSM requires a relatively large number of runs per factor and varies each factor over three to five levels. When selecting an experimental design, experimenters need to define the design space and ranges for each factor. Our full RSM design included three factors: total RNA mass, probe mass, and volume of hydrophillic streptavidin-coated magnetic bead solution. We require at least 100 ng of total RNA to prepare an RNA-seq library and used the level in the current protocol (400 ng) as an upper limit. To find a suitable range for the probes, we varied the concentration of each probe over five levels and searched for conditions that minimized the the average rRNA and probe abundance (Supplementary Table 1). Runs with the lowest overall probe concentration (1000–1400 ng combined) performed best. Given these results, our RSM design varied the probe mass between 200–800 ng per probe. We fixed the ratio of 16s and 23s probes at 1:1 by mass for all runs. The volume of beads must be no less than the combined volume of the probes and total RNA to ensure proper capture. The concentration of the probes and the well size of our 96-well plates constrain the bead volume to between 50 and 150 *μ*l.

We selected a rotatable central composite design (CCD) [13] with five levels for each factor (Figure 2B). The CCD provides optimal estimates of quadratic effects and interactions between factors. The final three-factor design includes 17 runs (Supplementary Table 2). We measured rRNA and probe abundance by qPCR as the response. Figure 3A shows the abundance of each run plotted in descending order of response. This run ordering confirmed our observations that lower levels of probe reduces rRNA and probe abundance.

**Figure 3:**
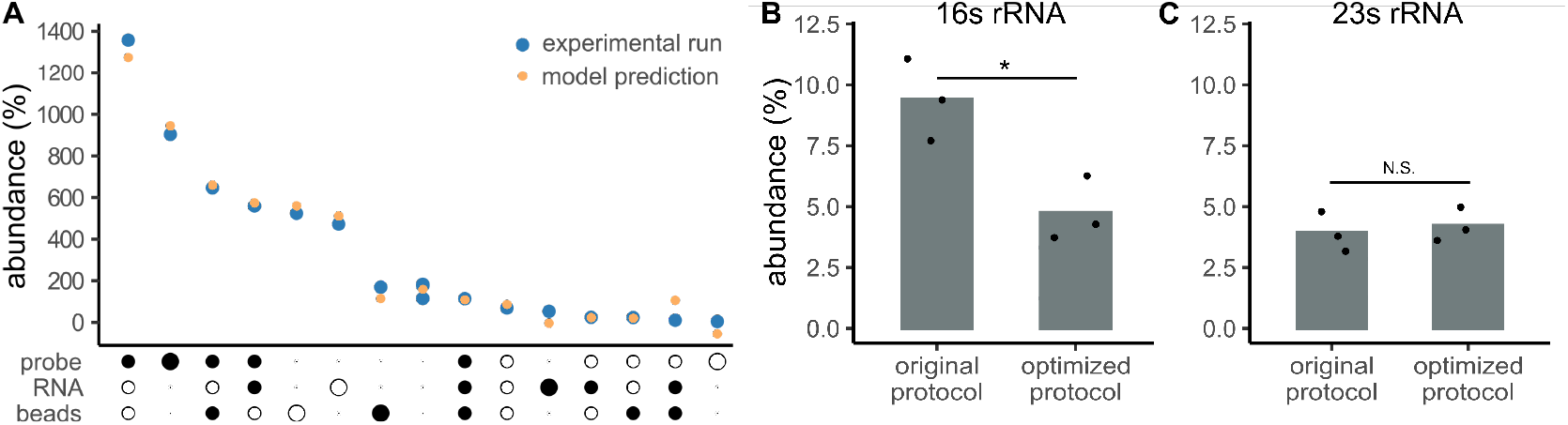
A second-order linear model predicts how rRNA depletion depends on the amount of probe, total RNA, and streptavadin beads. The response measures both rRNA and unbound probe remaining in the sample. **A**. A factor-and-response plot shows the rRNA and probe abundance (blue) along with the model prediction (orange) for each experimental run. Factor settings are encoded on the horizontal axis with their size proportional to their coded level. An unfilled circle indicates a negative coded value, while a filled circle indicates a positive coded value. **B**,**C**. The final optimized protocol removes more 16s rRNA (*p* = 0.02 by ANOVA) and requires fewer reagents. The optimized protocol removes up to 95% of the 16s and 23s rRNAs.

We fit a linear model to the response data to quantify the relationship between abundance and the three factors. The fitted model (showing only significant effects) is

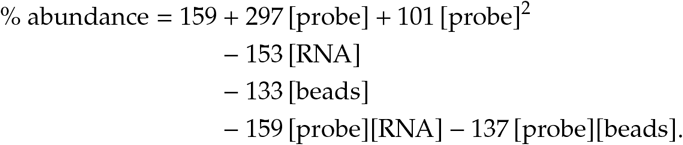

The model uses coded factors with levels − 1.68, − 1, 0, +1, and +1.68. The model is nonlinear with significant main effects, two way interactions, and pure quadratic terms (*p* < 0.05 by ANOVA), and no significant lack of fit (*p* > 0.19 by *F*-test). The model’s residual standard error is 69.9% and the adjusted *R*^2^ = 0.96.

As mentioned above, adding more probe increases the abundance of rRNA. Our qPCR assay measures both rRNA and probe; the increased abundance may be caused by leftover probe that is not removed by the beads. The significant quadratic effect creates a nonlinear relationship between probe concentration and rRNA depletion. For example, the model predicts that increasing the probe mass from 678 ng (the +1 level) to 800 ng (the +1.68 level) while holding other factors at the 0 level would increase the measured rRNA and probe abundance by 69%. We believe the nonlinear effects of adding probe are due to bead saturation because of the significant interaction between the probes and beads. Adding more beads is predicted to decrease the abundance of rRNA and probe while offsetting the detrimental effects of high probe levels.

The most interesting result from the model is the effect of total RNA. We expected that increasing total RNA would increase the percent abundance, but the model predicts the opposite—more RNA increases the relative depletion. There is a strong interaction between RNA and probe levels, making it difficult to interpret the effects of RNA directly. The levels of total RNA, probe, and beads must be considered simultaneously when optimizing the protocol. Changing any one of the levels without adjusting the others can easily overwhelm the probe’s ability to bind rRNA and the beads’ ability to remove all the probes.

We used numerical optimization to search for an improved protocol. A quadratic model can have a single optimum or a saddle point that indicates there is a tradeoff between different factors. Eigenanalysis of our model indicates a saddle point, so there is not a single best set of factor levels that maximize rRNA depletion. Moreover, the saddle point lies outside of the region explored by our experiments, and caution must be used when extrapolating so far outside the data used to fit the model. Instead, RSM practitioners recommend using a ridge analysis to follow the best direction calculated by the model and perform a second round of RSM at the new factor settings [14], [15].

Ridge analysis directed us toward the run with the lowest probe levels (200 ng). We began a second round of RSM centered at this new point by varying the amount of total RNA and the volume of beads. The CCD experimental design can be split into two blocks: the factorial runs and the axial runs. Experimenters can first run the factorial block and use it as a screen. Factors with small or insignificant effect sizes can be removed from the later axial block to reduce the number of runs. The factorial block of our CCD indicated that runs at the center point performed best, so there was no need to continue with the axial block (Supplementary Table 3). Increasing the total RNA or reducing the volume of beads increased the percent abundance, and increasing the volume of beads did not improve depletion efficiency.

Our percent abundance measurement represents the average of the 16s and 23s rRNAs. In experiments where probes were added in equal mass, we noticed that the 16s probes outperformed the 23s probes (< 5% vs. 10%). We hypothesized that since the lower molecular weight 16s probe was in a higher *molar* abundance, the probes were better able to capture the 16s rRNA. To compensate for this difference, we increased the mass of the 23s probe from 200 ng to 250 ng. Indeed, this improved 23s rRNA capture without over-saturating the beads.

The final, optimal conditions for 16s and 23s rRNA depletion appear in Table 1. Compared to the original protocol in Stewart, *et al*, our optimized protocol is more efficient at removing rRNA (4.7% vs. 9.4% abundance for 16s, 4.2% vs. 3.9% for 23s) (Figure 3B,C). We confirmed the rRNA depletion in our optimized protocol with capillary electrophoresis (Supplementary Figure 1).

**Table 1:**
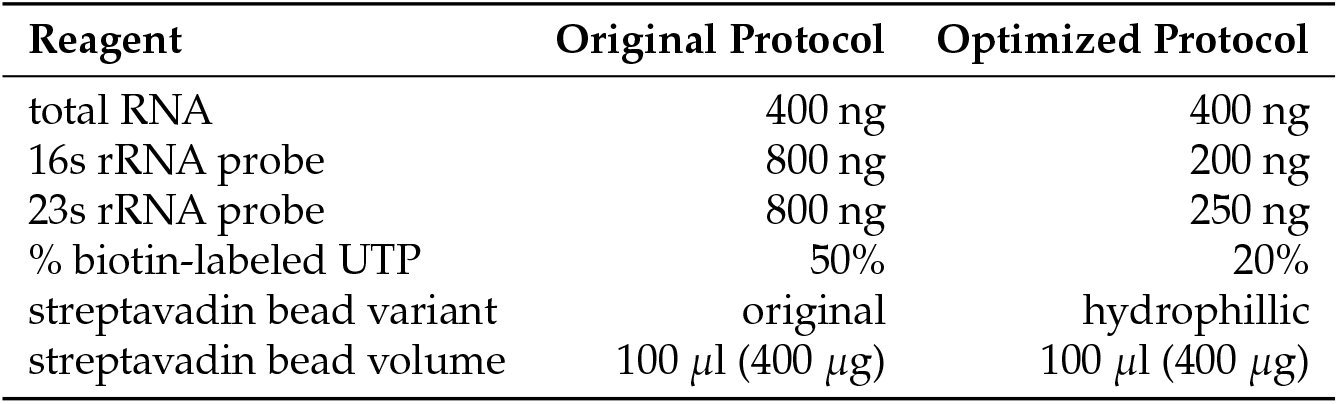
Original and optimzied rRNA depletion reagent settings.

| Reagent | Original Protocol | Optimized Protocol |
| --- | --- | --- |
| total RNA | 400 ng | 400 ng |
| 16s rRNA probe | 800 ng | 200 ng |
| 23s rRNA probe | 800 ng | 250 ng |
| % biotin-labeled UTP | 50% | 20% |
| streptavidin bead variant | original | hydrophilic |
| streptavidin bead volume | 100 μl (400 μg) | 100 μl (400 μg) |

### 2.3 Cost analysis

The reagent costs for our optimized rRNA depletion assay appear in Supplementary Material 4. The streptavidin-coated magnetic beads are the largest single expense, accounting for 63% of the total reagent cost. We tested if the volume of beads could be reduced without sacrificing the rRNA depletion efficiency. As reported in Supplementary Figure 2, the volume of beads can be reduced from 100 *μ*l to 75 *μ*l without loss of efficiency. Unless reducing costs is a primary concern, we recommend using 100 ul to ensure full removal of rRNA and the probes. We also used the excess of beads when adding antisense probes for the 5s rRNA, as discussed in the following section. Even with 100 *μ*l of beads, our optimized protocol costs 14.7% less than the original protocol.

### 2.4 Designing probes for the 5s rRNA

The original protocol in Stewart, *et al*. includes probes for the 16s and 23s rRNAs, but not the 5s rRNA. We created antisense RNA probes for the 5s rRNA and measured depletion by qPCR. Initially, the depletion of 5s rRNA was poor (60–80% abundance), implying either poor hybridization to the rRNA or probes left uncaptured to the beads. We suspected that the 27 uracils in the 116 bp 5s rRNA created too few sites for potential biotin labeling. Indeed, when using 20% labeled UTPs (the level used for the 16s and 23s probes) approximately 7% of the probes would have less than three biotin modifications. Increasing the concentration of labeled uracils on the 5s rRNA probes from 20% to 50% improved the depletion of the 5s rRNA to levels similar to the 16s and 23s rRNAs.

The final protocol for rRNA depletion with 5s probes includes 200 ng 16s probe, 250 ng 23s probe, 50 ng 5s probe, 400 ng total RNA, and 100 *μ*l beads. A step-by-step protocol is available in Supplementary Material 5.

## 3 Discussion

This work used statistical design of experiments to optimize an rRNA depletion protocol. Protocols require optimization for two reasons. First, reagents can change over time. For example, the streptavadin beads used by Stewart, *et al*. have a higher binding capacity for both labeled and unlabeled nucleic acid than the hydrophillic streptavadin beads used in our protocol. Switching to the hydrophillic beads required changes in probe levels, biotin concentration, and total RNA in the reaction. Second, the original protocol developers may not have fully explored the design space. Some unexplored factors or unknown interactions may impact protocol performance. Conversely, other factors may be expensive but not have a large impact on performance. For example, lowering the concentration of biotin on the probes did not improve rRNA depletion, but it lowered the cost of probe synthesis by over 50%.

RSM identifies interactions that are missed by one-at-a-time optimization. Interactions make it difficult to predict the relationship between factors the response. For example, the optimal concentration of probe depends on the amount of beads, and we cannot explain the beneficial interaction between probe and RNA concentration. Our quantitative model allowed us to find improved conditions despite the nonlinear interactions between factors.

Small increases in rRNA depletion efficiency can significantly increase mRNA abundanace in sequencing libraries. Because rRNAs represent >80% of the total RNA in a bacterial cell, they dominate sequencing libraries unless they are depleted. Our optimized protocol removes >95% of the rRNA, so non-rRNA transcipts make up 85% of the remaining transcripts. There is still room for improvement. If rRNA depletion efficiency increased to 97.5%, non-rRNA transcipts would make up 92% of the sample, an 8% increase over our optimized protocol. Additionally, one could consider designing probes that target tRNAs or other highly abundant transcripts to further increase the proportion of mRNA in the library. Any improvements to mRNA enrichment improve the depth and lower the cost of downstream sequencing.

Users who design probes for rRNAs or other transcripts may need to empirically determine the optimal ratio of labeled to unlabeled nucleotides. It is the total number of biotin modifications in a probe, not the ratio of labeled to unlabeled nucleotides, that matters. Our experiments indicated that for large transcripts such as the 16s and 23s rRNAs, a large excess of biotin labels unnecessarily increased cost. Conversely, having too few modifications resulted in inefficient capture, as was the case for our original 5s rRNA probes.

DOE and RSM can optimize protocols efficiently, with few runs per factor. These techniques allow protocol developers to consider more factors and produce better protocols. DOE and RSM can also quantify the relationship between factors and the response, which is helpful when making rational, data-driven decisions about design tradeoffs. We hope this work increases the use of DOE and RSM in molecular and synthetic biology.

## 4 Methods

### 4.1 RNA and DNA extraction

*Streptococcus mutans* strain UA159 was grown in THY (Todd Hewitt Broth + 0.5% yeast extract, Millipore-Sigma) at 37 °C and 5% CO_2_ and harvested during mid-log phase growth. DNA was extracted using the DNeasy® UltraClean Microbial Kit (QIAGEN) and stored at − 20 °C. For RNA extraction, cultures were centrifuged, aspirated, and re-suspended in TRIZOL reagent (Ambion) and ≈ 200 *μL* of 0.1 mm diameter silica beads (BioSpec). Samples were homogenized for 3 minutes in two 90 s intervals at 1600 rpm (OHAUS homogenizer), then heated at 65 °C for 5 minutes. RNA extraction and purification was performed using the DirectZol® RNA Minprep Plus Kit (Zymo). Extracted RNA was stored at −80 °C.

### 4.2 Probe design and synthesis

A detailed protocol for probe design and rRNA depletion is provided in Supplementary Material 5. Primers (IDT) for amplifying probe templates targeted the *S. mutans* rRNA genes (Supplementary Table 4) and included a T7 promoter sequence at 5’ end of the reverse primer. The DNA templates were PCR amplified using Q5 DNA Polymerase (NEB), purified using the GeneJet PCR Purification Kit (Thermo), and size verified by gel electrophoresis. *In vitro* transcription (IVT) was performed with the HiScribe™ T7 RNA Synthesis Kit (NEB). For probes in the original protocol, the biotin-labeled UTP concentration was 50% (Supplementary Material 3). The optimized probes contained 20% biotin-labeled UTP for the 16s and 23s probes and 50% for the 5s rRNA. Following IVT, the probes were purified with the Monarch® RNA Cleanup Kit (500 *μ*g, NEB).

### 4.3 rRNA depletion

rRNA probes were hybridized to total RNA at 70 °C for 5 minutes, followed by a ramp down to 25 °C by 5 °C increments for 1 minute each. Meanwhile, regular (NEB S1420S) or hydrophillic (NEB S1421S) streptavadin magnetic beads were aliqoted and washed in 0.1 N NaOH followed by two washes in 1× SSC buffer (Invitrogen). Hybridized RNA and probes were diluted to the original bead volume using a 1 × SSC and 20% formamide solution and incubated at room temperature for 3 minutes prior to bead capture. RNA and probe hybrids were bound to the washed beads for 10 minutes at room temperature and separated using a magnetic rack. The rRNA depleted supernatant was purified with the Monarch®RNA Cleanup Kit (10*μ*g) before downstream analysis.

### 4.4 qPCR assay for rRNA depletion

qPCR primers (IDT) were designed to amplify both the leftover probes and rRNA (Supplementary Table 5). The *S. mutans* housekeeping gene *ldh* was amplified as a baseline signal control. PCRs were performed using the Luna® Universal One Step RT-qPCR Kit (NEB) in a ThermoFisher Quantstudio 3 instrument. Cycle threshold (C_t_) values were measured for the rRNA targets and ldh gene in both depleted and undepleted samples. rRNA and probe abundance was calculated using the equation

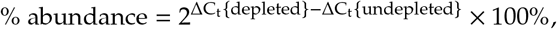

where

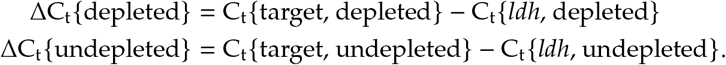

The average abundance of the 16s and 23s rRNAs was used as the response for our RSM experiments.

### 4.5 Statistical analysis

Statistical analysis was performed using R version 3.6.2 with the rsm package [16]. We created a model for rRNA abundance that included first order, two-way interaction, and pure quadratic terms. Factor significance was assessed by ANOVA and *t*-test.

## 5 Acknowledgements

We thank Bill Metcalf for introducing us to the original rRNA depletion protocol and Walden Li for independently testing the optimized protocol. This work was supported by the National Institutes of Health grant GM138210. BMD is supported in part by an Illinois Distinguished Fellowship.

