## Supplementary Material for "Improving an rRNA depletion protocol with statistical design of experiments"

### 1 Supplementary Figures

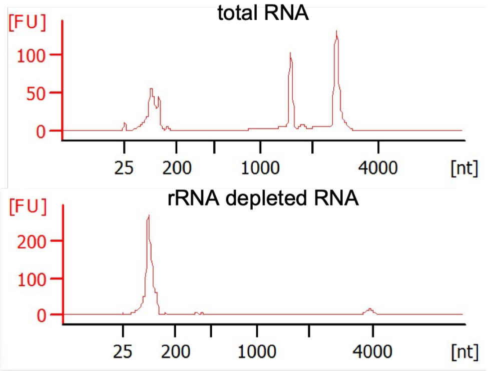


**Supplementary Figure 1:** Capillary electrophoresis confirms rRNA removal using the optimized protocol. The 1557 bp 16s rRNA and 2910 bp 23s rRNA are not detectable in the rRNA depleted samples. Samples were loaded at an equal mass.

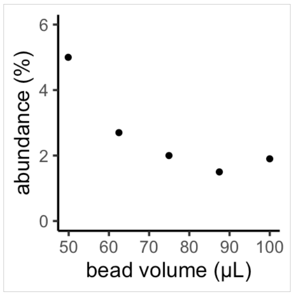


**Supplementary Figure 2:** Bead volumes between 75–100 $\mu$l efficiently remove the 16s and 23s rRNA. We recommend using 100 $\mu$l to ensure efficient removal unless reducing cost is the primary concern.

### 2 Supplementary Tables

**Supplementary Table 1:** A 2 factor Central Composite Design for individual probe levels. A CCD with 4 factorial points is rotatable when $\alpha = \sqrt[4]{4} \approx 1.414$.

| Run | 16s Probe | 23s Probe | 16s Probe ($\mu$g) | 23s Probe ($\mu$g) | Abundance (%) |
|---|---|---|---|---|---|
| 1 | 0 | 0 | 1000 | 1000 | 119 |
| 2 | 0 | 0 | 1000 | 1000 | 128 |
| 3 | 0 | 0 | 1000 | 1000 | 142 |
| 4 | 0 | 0 | 1000 | 1000 | 162 |
| 5 | 0 | 0 | 1000 | 1000 | 97.2 |
| 6 | −1 | −1 | 576 | 576 | 68.7 |
| 7 | 1 | −1 | 1424 | 576 | 160 |
| 8 | −1 | 1 | 576 | 1424 | 131 |
| 9 | 1 | 1 | 1424 | 1424 | 406 |
| 10 | $-\alpha$ | 0 | 400 | 1000 | 51.4 |
| 11 | $\alpha$ | 0 | 1600 | 1000 | 420 |
| 12 | 0 | $-\alpha$ | 1000 | 400 | 109 |
| 13 | 0 | $\alpha$ | 1000 | 1600 | 287 |

**Supplementary Table 2:** A 3 factor Central Composite Design to optimize probe, RNA, and bead levels. A CCD with 8 factorial points is rotatable when $\alpha = \sqrt[4]{8} \approx 1.682$.

| Run | Probes | RNA | Beads | Probes ($\mu$g) | RNA ($\mu$g) | Beads ($\mu$l) | Abundance (%) |
|---|---|---|---|---|---|---|---|
| 1 | −1 | −1 | −1 | 322 | 161 | 70 | 69.2 |
| 2 | 1 | −1 | −1 | 678 | 161 | 70 | 1360 |
| 3 | −1 | 1 | −1 | 322 | 339 | 70 | 24.4 |
| 4 | 1 | 1 | −1 | 678 | 339 | 70 | 560 |
| 5 | −1 | −1 | 1 | 322 | 161 | 130 | 23.4 |
| 6 | 1 | −1 | 1 | 678 | 161 | 130 | 647 |
| 7 | −1 | 1 | 1 | 322 | 339 | 130 | 10.0 |
| 8 | 1 | 1 | 1 | 678 | 339 | 130 | 114 |
| 9 | 0 | 0 | 0 | 500 | 250 | 100 | 115 |
| 10 | 0 | 0 | 0 | 500 | 250 | 100 | 183 |
| 11 | 0 | 0 | 0 | 500 | 250 | 100 | 176 |
| 12 | $-\alpha$ | 0 | 0 | 200 | 250 | 100 | 4.31 |
| 13 | $\alpha$ | 0 | 0 | 800 | 250 | 100 | 904 |
| 14 | 0 | $-\alpha$ | 0 | 500 | 100 | 100 | 472 |
| 15 | 0 | $\alpha$ | 0 | 500 | 400 | 100 | 53.2 |
| 16 | 0 | 0 | $-\alpha$ | 500 | 250 | 50 | 524 |
| 17 | 0 | 0 | $\alpha$ | 500 | 250 | 150 | 169 |

**Supplementary Table 3:** A $2^2$ Factorial Design with center points at the optimum found in Supplementary Table 2.

| Run | Probes | RNA | Beads | Probes ($\mu$g) | RNA ($\mu$g) | Beads ($\mu$l) | Abundance (%) |
|---|---|---|---|---|---|---|---|
| 1 | 0 | 0 | 0 | 200 | 400 | 100 | 1.13 |
| 2 | 0 | 0 | 0 | 200 | 400 | 100 | 2.14 |
| 3 | 0 | −1 | −1 | 200 | 250 | 50 | 5.20 |
| 4 | 0 | 1 | −1 | 200 | 550 | 50 | 16.9 |
| 5 | 0 | −1 | 1 | 200 | 250 | 150 | 1.36 |
| 6 | 0 | 1 | 1 | 200 | 550 | 150 | 6.96 |

**Supplementary Table 4:** Primers for amplifying probe templates in *S. mutans* UA159. The T7 promoter sequence is underlined. Annealing temperatures and extension times are for the Q5 polymerase (NEB).

| Target | Sequence |
|---|---|
| **16s rRNA** | 1552 bp amplicon, $T_a$ = 65 °C, 45 s extension time |
| forward | AGAGTTTGATCCTGGCTCAG |
| reverse | <u>GCCAGTGAATTGTAATACGACTCACTATAGGG</u>ACGGCTACCTTGTTACGACTT |
| **23s rRNA** | 2325 bp amplicon, $T_a$ = 56 °C, 75 s extension time |
| forward | GAACTGAAACATCTCAGTA |
| reverse | <u>GCCAGTGAATTGTAATACGACTCACTATAGGG</u>CGACATCGAGGTGCCAAA |
| **5s rRNA** | 120 bp amplicon, $T_a$ = 58 °C, 25 s extension time |
| forward | TTAAGTGATGATAGCCTAGG |
| reverse | <u>GCCAGTGAATTGTAATACGACTCACTATAGGG</u>TCTTGCTAAGCGACGA |

**Supplementary Table 5:** qPCR primers to quantify depletion efficiency in *S. mutans* UA159. Primers target the middle of each gene. The *ldh* gene (SMU_1115) was used as a control.

| Target | Sequence |
|---|---|
| **16s rRNA** | 85 bp amplicon, $T_m$= 61.5 °C |
| forward | TCGAAAGCGTGGGTAGCGAACA |
| reverse | CCGGAAAGGGCCTAACACCTAGC |
| **23s rRNA** | 75 bp amplicon, $T_m$= 61 °C |
| forward | TGAGTGAAGGAGGGACGCAGCA |
| reverse | GTCCTCACCTCACTGTTGGACG |
| **5s rRNA** | 75 bp amplicon, $T_m$= 60 °C |
| forward | TGCCGAACACAGCAGTTAAGCCC |
| reverse | AGCGACGACCCTATCTCACAGG |
| ***ldh*** | 127 bp amplicon, $T_m$= 61 °C |
| forward | TCCTCGTTGCTGCTAACCCAGT |
| reverse | GCAAGTGCTTGACGGAAACGAGC |

### 3 Biotin-labeling Distributions

The number of biotins per molecule of probe can be modeled as a set of independent binomial processes, one for each type of nucleotide (U, C, G, or A). The number of labeled uracils is a random variable

$$U \sim \text{Binomial}(n_U, p_U)$$

where $n_U$ is the total number of uracils and $p_U$ is the probability of incorporating a biotin-labeled UTP in each of the $n_U$ sites. During *in vitro* transcription, the probability of adding a biotin-labeled nucleotide is equal to the fraction of biotin-labeled nucleotides in the reaction mixture. The original Stewart, *et al.* protocol included 25% biotin-labeled CTP and UTP and no biotin-labled ATP or GTP. Under the binomial assumption, the expected value and variance for the number of labeled uracils is

$$\begin{aligned} \mathrm{E}[U] &= n_U p_U \\ \mathrm{Var}[U] &= n_U p_U (1 - p_U), \end{aligned}$$

and similar statistics can be calculated for the number of labeled cytosines ($C \sim \text{Binomial}(n_C, p_C)$).

The binomial processes for biotin labeling of separate nucleotides are assumed to be independent, so the total number of biotins per probe ($X$) is

$$\begin{aligned} \mathrm{E}[X] &= \mathrm{E}[U] + \mathrm{E}[C] \\ &= n_U p_U + n_U p_U \\ \mathrm{Var}[X] &= \mathrm{Var}[U] + \mathrm{Var}[C] \\ &= n_U p_U (1 - p_U) + n_U p_U (1 - p_U). \end{aligned}$$

The original Stewart, *et al.* probes had equal fractions of labeled uracil and cytosine ($p_U = p_U = \lambda = 0.25$). The expected value and variance for the original protocol are

$$\begin{aligned} \mathrm{E}[X] &= n_U p_U + n_C p_C \\ &= (n_U + n_U)\lambda \\ \mathrm{Var}[X] &= n_U p_U (1 - p_U) + n_U p_U (1 - p_U) \\ &= (n_U + n_U)\lambda(1 - \lambda). \end{aligned}$$

We approximated the Stewart, *et al.* probes by removing the biotin-labeled CTPs and doubling the fraction of labled UTP to 50% ($p_U = 0$ and $p_U = 2\lambda = 0.5$). The expected number of biotins and the associated variance for our probes are

$$\begin{aligned} \mathrm{E}[X] &= 2 n_U \lambda \\ \mathrm{Var}[X] &= 2 n_U \lambda (1 - 2\lambda). \end{aligned}$$

The *S. mutans* 16s and 23s rRNA probes have similar numbers of uracils and cytosines ($n_U = 400$ and 807, $n_C = 476$ and 895 respectively), so labeling only uracil (at twice the rate) does not change the expected number of biotins per probe. Using only labeled uracils reduces the reagent costs and reduces the variance in the number of biotins per molecule.

$$\begin{aligned} \mathrm{Var}[X_{U\&C}] &= (n_U + n_C)\lambda(1 - \lambda) \\ &\approx 2 n_U \lambda (1 - \lambda) \\ &= 0.375 n_U \\ \mathrm{Var}[X_{U \text{ only}}] &= 2 n_U \lambda (1 - 2\lambda) \\ &= 0.125 n_U. \end{aligned}$$

Our optimized protocol uses only labeled uracils and decreases the biotin concentration from 50% to 20% ($p_U = 0.2$). The expected number of biotins on the optimized probes is

$$\mathrm{E}[X] = 0.2 n_U$$

with variance

$$\mathrm{Var}[X] = 0.2 n_U (1 - 0.2) = 0.16 n_U.$$

We simulated biotin incorporation for each of the probe sets. The results of the simulation and a comparison to the theoretical calculations appear in Supplementary Figure 3 and in Supplementary Table 6.

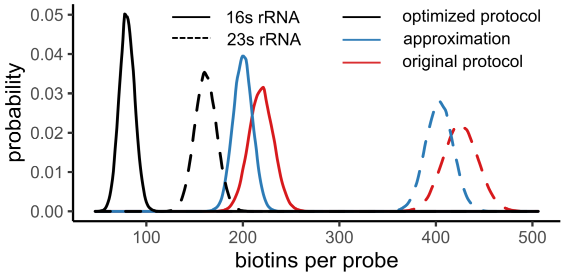


**Supplementary Figure 3:** The optimized protocol decreases the overall biotin concentration, reducing the cost of probe synthesis by over 50%

**Supplementary Table 6:** Theoretical calculations of biotin-labling distributions for antisense rRNA probes.

| | | Original protocol | Approximation | Optimized protocol |
|---|---|---|---|---|
| **16s rRNA** | E[X] | 219.0 | 200.0 | 80.0 |
| | Var[X] | 164.3 | 100.0 | 64.0 |
| **23s rRNA** | E[X] | 425.5 | 403.5 | 161.4 |
| | Var[X] | 319.1 | 201.8 | 129.1 |

### 4 Cost Analysis

#### 4.1 PCR amplification of rRNA genes

**Cost to amplify each rRNA gene**

| Reagent | Original Protocol | Optimized Protocol |
|---|---|---|
| Q5 Hot-Start High-Fidelity DNA Polymerase Mix | $2.17 | $2.17 |
| GeneJET PCR Purification Kit | $1.70 | $1.70 |
| **Total Cost** | **$3.87** | **$3.87** |

#### 4.2 IVT Probe Synthesis

**Cost per *in vitro* transcription reaction**

| Reagent | Original Protocol | Optimized Protocol |
|---|---|---|
| HiScribe$^{TM}$ T7 High Yield RNA Synthesis Kit | $4.64 | $4.64 |
| Bio-16 UTP | $45.30 | $36.24 |
| Bio-11 CTP | $48.90 | ——— |
| rRNA Gene PCR Products | $0.15 | $0.15 |
| **Total Cost** | **$98.99** | **$41.03** |

##### 4.3 rRNA Depletion

**Cost per sample of total RNA**

| Reagent | Original Protocol | Optimized Protocol |
|---|---|---|
| Streptavadin Magnetic Beads | $6.68 | $6.68 |
| 0.1 N NaOH | < $0.01 | < $0.01 |
| 20× SSC Buffer | $0.05 | $0.05 |
| RNase Inhibitor, Murine | $0.77 | $0.77 |
| Formamide | < $0.01 | < $0.01 |
| Monarch RNA Cleanup Kit (10 $\mu$g) | $2.90 | $2.90 |
| 16s and 23s Probes | $2.11 | $0.27 |
| **Total Cost** | **$12.51** | **$10.67** |

### 5 Optimized Protocol

#### Reagents

| Reagent | Supplier | Product # |
|---|---|---|
| Q5 Hot Start DNA Polymerase Mix (2×) | NEB | M0494 |
| GeneJet PCR Purification Kit | Thermo | K0701 |
| HiScribe T7 High Yield RNA Synthesis Kit | NEB | E2040 |
| Bio-16 UTP (10 mM) | Thermo | AM8452 |
| Monarch RNA Cleanup Kit (500 $\mu$g) | NEB | T2050 |
| Hydrophillic Streptavadin Magnetic Beads | NEB | S1421 |
| 0.1N NaOH | VWR (Amresco) | E584 |
| 20× SSC Buffer | Thermo | AM9770 |
| RNase Inhibitor, Murine | NEB | M0134 |
| Formamide (100%) | Thermo (Alfa Aesar) | A11076 |
| Monarch RNA Cleanup Kit (10 $\mu$g) | NEB | T2030 |
| Luna Universal One Step RT-qPCR Kit (optional) | NEB | E3005 |

#### 5.1 PCR Amplify rRNA Genes

1. Design primers for antisense rRNA templates. Primers should amplify the full length rRNA gene. Include the T7 promoter sequence (GCCAGTGAATTGTAATACGACTCACTATAGGG) on the 5′ end of the reverse primer.

2. Prepare PCRs for the rRNA probe templates

| Reagent | 50 $\mu$l reaction | Final Concentration |
|---|---|---|
| Q5 HF 2X Master Mix | 25 $\mu$l | 1× |
| Forward Primer (10 $\mu$M) | 2.5 $\mu$l | 0.5 $\mu$M |
| Reverse Primer (10 $\mu$M) | 2.5 $\mu$l | 0.5 $\mu$M |
| gDNA Template | X $\mu$l | 50 ng |
| Nuclease-Free Water | to 50 $\mu$l | |

**PCR Cycling Conditions**

| Step | Temperature | Time |
|---|---|---|
| Initial Denaturation | 98 °C | 30 s |
| 30 cycles | 98 °C<br>X °C (use NEB $T_m$ calculator)<br>X 72 °C | 10 s<br>20 s<br>X s (≈ 30 s/kb) |
| Final Extension | 72 °C | 120 s |

3. Purifiy PCR products using the GeneJet PCR purification kit according to the manufacturer protocol. Elute the PCR products in 30 $\mu$l nuclease-free water. Verify the products by gel electrophoresis before proceeding.

#### 5.2 *in vitro* transcription of biotinylated RNA probes

1. Thaw all components of the HiScribe T7 High Yield RNA Synthesis Kit, keep all components on ice.

2. Assemble the following IVT reactions at room temperature. Note the different biotin concentration for the 5s rRNA probes.

| Reagent | 16s/23s probes | Concentration | 5s probes | Concentration |
|---|---|---|---|---|
| 10× Reaction Buffer | 1.5 $\mu$l | 0.75× | 1.5 $\mu$l | 0.75× |
| ATP (100 mM) | 1.5 $\mu$l | 7.5 mM | 1.5 $\mu$l | 7.5 mM |
| GTP (100 mM) | 1.5 $\mu$l | 7.5 mM | 1.5 $\mu$l | 7.5 mM |
| CTP (100 mM) | 1.5 $\mu$l | 7.5 mM | 1.5 $\mu$l | 7.5 mM |
| UTP (100 mM) | 1.2 $\mu$l | 6 mM | 0.75 $\mu$l | 3.75 mM |
| Bio-16 UTP (10 mM) | 3 $\mu$l | 1.5 mM | 7.5 $\mu$l | 3.75 mM |
| Template DNA | X $\mu$l | 100 ng | X $\mu$l | 100 ng |
| T7 RNA Polymerase Mix | 1.5 $\mu$l | | 1.5 $\mu$l | |
| Nuclease-free water | to 20 $\mu$l | | to 20 $\mu$l | |

3. Mix thoroughly by pipetting, pulse spin, and incubate at 37 °C for 2 hours.
4. Purify the IVT products with the Monarch RNA cleanup kit (500 $\mu$g). Elute in 50 $\mu$l nuclease-free water.
5. **Optional Quality Control**: verify product size and purity by gel electrophoresis.

##### 5.3 rRNA Depletion

1. Bead Washing (can be performed before or during probe hybridization)
   (a) Aliquot 100 $\mu$l beads per sample into individual microcentrifuge tubes.
   (b) Bind beads to a magnetic tube rack (≈2 minutes), aspirate and discard the supernatant, and resuspend in 100 $\mu$l 0.1N NaOH to deactivate bead-associated RNases. Mix well by pipetting, re-capture the beads on the magnetic rack, and aspirate and discard the supernatant
   (c) Wash the beads twice more using 1× SSC buffer. After the second wash, do not aspirate the supernatant. Instead, leave the beads suspended and on ice until the hybridization reaction is complete.
2. Prepare the following hybridization reactions for each sample.

| Reagent | Volume |
|---|---|
| total RNA template (400 ng) | X $\mu$l |
| 16s rRNA probe (200 ng) | X $\mu$l |
| 23s rRNA probe (250 ng) | X $\mu$l |
| 5s rRNA probe (50 ng) (optional) | X $\mu$l |
| RNase inhibitor, murine | 1 $\mu$l |
| 20× SSC buffer | 2.5 $\mu$l |
| Formamide (100%) | 10 $\mu$l |
| Nuclease-free water | to 50 $\mu$l |

   (a) Incubate the hybridization reactions at 70 °C for 5 minutes follwed by a ramp down to 25 °C at 5 °C increments of 1 minute each.
   (b) After hybridization, incubate the reactions at room temperature for 5 minutes.
3. Bead Binding
   (a) While the hybridization reactions are incubating at room temperature, capture the cleaned beads on the magnetic rack and aspirate the supernatant. Remove the beads from the rack
   (b) Dilute the hybridization reactions up to 100 $\mu$l with a 1X SSC buffer and 20% formamide solution (20 $\mu$l formamide for every 80 $\mu$l of 1X SSC buffer).
   (c) Add the hybridization reactions to the cleaned beads

(d) Incubate the solution at room temperature for 10 minutes. Mix the samples twice by pipetting during the incubation.

4. Capture the beads on the magnetic rack. Transfer the rRNA depleted supernatant to an RNase-free tube. Discard the used beads.

5. Purify the rRNA depleted RNA using the Monarch RNA cleanup kit (10 $\mu$g). Elute in 20 $\mu$l nuclease-free water. Store the RNA at −80 °C for downstream applications

6. *Optional Quality Control Assay*. Verify the rRNA depletion efficiency either by qPCR or bioanalyzer
